# Age and mortality associated DNA methylation patterns on the X-chromosome in male and female samples

**DOI:** 10.1101/636597

**Authors:** Shuxia Li, Jesper B. Lund, Jan Baumbach, Kaare Christensen, Jonas Mengel-From, Weilong Li, Afsaneh Mohammadnejad, Alison Pattie, Riccardo E. Marioni, Ian J. Deary, Qihua Tan

**Affiliations:** Unit of Human Genetics, Department of Clinical Research, University of Southern Denmark, Odense, Denmark; Epidemiology and Biostatistics, Department of Public Health, University of Southern Denmark, Odense, Denmark; Chair of Experimental Bioinformatics, TUM School of Life Sciences Weihenstephan, Technical University of Munich, Germany; Department of Psychology, University of Edinburgh, Edinburgh, Scotland, United Kingdom; Centre for Genomic and Experimental Medicine, University of Edinburgh, Edinburgh, Scotland, United Kingdom; Centre for Cognitive Aging and Cognitive Epidemiology, University of Edinburgh, Edinburgh, Scotland, United Kingdom

**Keywords:** X-chromosome, DNA methylation, age, mortality

## Abstract

**Background:** Multiple epigenetic association studies on human aging have been performed reporting large numbers of sites differentially methylated across ages on the autosomal chromosomes. The X-chromosome has been studied little, due to analytical difficulties in dealing with sex differences in X-chromosome content and X-inactivation in females. Based on large collections of genome-wide DNA methylation data on two Danish cohorts of identical twins (mean ages, 66 and 79 years) and the Lothian Birth Cohort 1921 (mean age 79 years), we conducted a chromosome-wide association analysis on male and female samples separately with equal sample sizes to discover age-dependent X-linked DNA methylation patterns using chromosome 20 with about same number of CpGs analysed as an autosomal reference, and compare the age-related changes in DNA methylation between the two sexes. In addition, age-related methylation sites were assessed for their associations with mortality.

**Results:** We identified more age-related DNA methylation sites (FDR<0.05) in females than in males. Among them, predominantly more sites were hypermethylated in the older as compared with the younger cohorts, a pattern similar to that observed on chromosome 20. Among the age-related sites, 13 CpGs in males and 24 CpGs in females were found significant (FDR<0.05) in all cohorts. Survival analysis showed that there are more age-methylated CpGs that contribute to reduce mortality than those that increase mortality in male but not in female samples.

**Conclusion:** The X-chromosome displays significant age-and sex-dependent methylation patterns which might be differentially associated with mortality in the two sexes.

## Introduction

Aging-related epigenetic changes have been studied intensively using high-throughput techniques for genome-wide DNA methylation profiling. There are reports of large numbers of significant CpG sites dominated by lower methylation with older age (Bell et al. 2012; Horvath et al. 2012; Hannum et al. 2013; Florath et al. 2013; Marttila et al. 2015; Tan et al. 2016; Moore et al. 2016). The age-associated methylation patterns have been recently replicated in different samples and even across populations (Li et al. 2017). Overall, findings from these studies point to the extensive epigenetic remodeling in the DNA methylome involving biological pathways related to aging phenotypes and age-related diseases. The X chromosome comprises about 5 percent of the human genome and harbours about 2,000 genes; however, methylation levels on this chromosome have typically been ignored to date. This is probably due to analytical difficulties in dealing with differences in X-chromosome contents of females and males, and X-chromosome inactivation in females which renders the comparison on levels of DNA methylation between the two sexes impossible.

Using a large collection of DNA methylation data on the Lothian Birth Cohort of 1921 and 2 Danish twin cohorts, we examine DNA methylation on the X-chromosome as a function of age, we compare it between male and female samples, and we assess their associations with mortality. The analysis took advantage of large sample sizes that enabled statistical modeling on male and female samples independently after a data normalization procedure conducted on male and female samples separately and on the X-chromosome only. Our strategic analysis simply focusing on the age-dependent change (or rate of change in a longitudinal setting) in DNA methylation helped us to circumvent X-chromosome complexity. The use of Danish twins helped to control for genetic influences on the aging process leading to enriched power for epigenetic association analysis (Li et al. 2018). Based on multiple comparison and replication, we show that, through both sex-specific and non-sex-specific age-related DNA methylation changes, the X-chromosome plays a non-neglectable role during the aging process.

## Results

By applying the linear regression models on male and female samples separately in three cohorts (Table 1), we identified X-linked CpGs displaying significant age-associated methylation changes (FDR<0.05) independent or dependent of sex with very high proportions of hypermethylation (≥66%) except for male-only CpGs from MADT (39%) (Table 2). In addition, higher proportions of hypermethylation are observed in the older LBC and LSADT cohorts than in the relatively younger MADT samples except for female-only CpGs from LSADT. Although equal numbers of male and female samples were used in the statistical analysis, far more significant sites were detected in female than in male samples with the smallest numbers of CpGs significant in both sexes (Table 2). By plotting the regression coefficient of age (i.e. the rate of change in DNA methylation across ages) in males against that in females, Figure 1 presents the age-methylated CpGs that are significant only in males (blue), in females (red) or in both sexes (green) in the 3 cohorts respectively. Comparing Figures 1a and 1c with 1e, one can see that, in general, the significant CpGs are more dominated by hypermethylation in the older cohorts than in the younger MADT cohort. A similar pattern was observed when plotting the estimates on chromosome 20 which has about the same number of CpG probes tested as chromosome X (Figures 1b, 1d and 1f). Comparing Figure 1a with 1b, one can see that the age-dependent methylation changes on chromosome X in the oldest LBC cohort is more diverse or heterogeneous then that on chromosome 20.

**Table 1.**
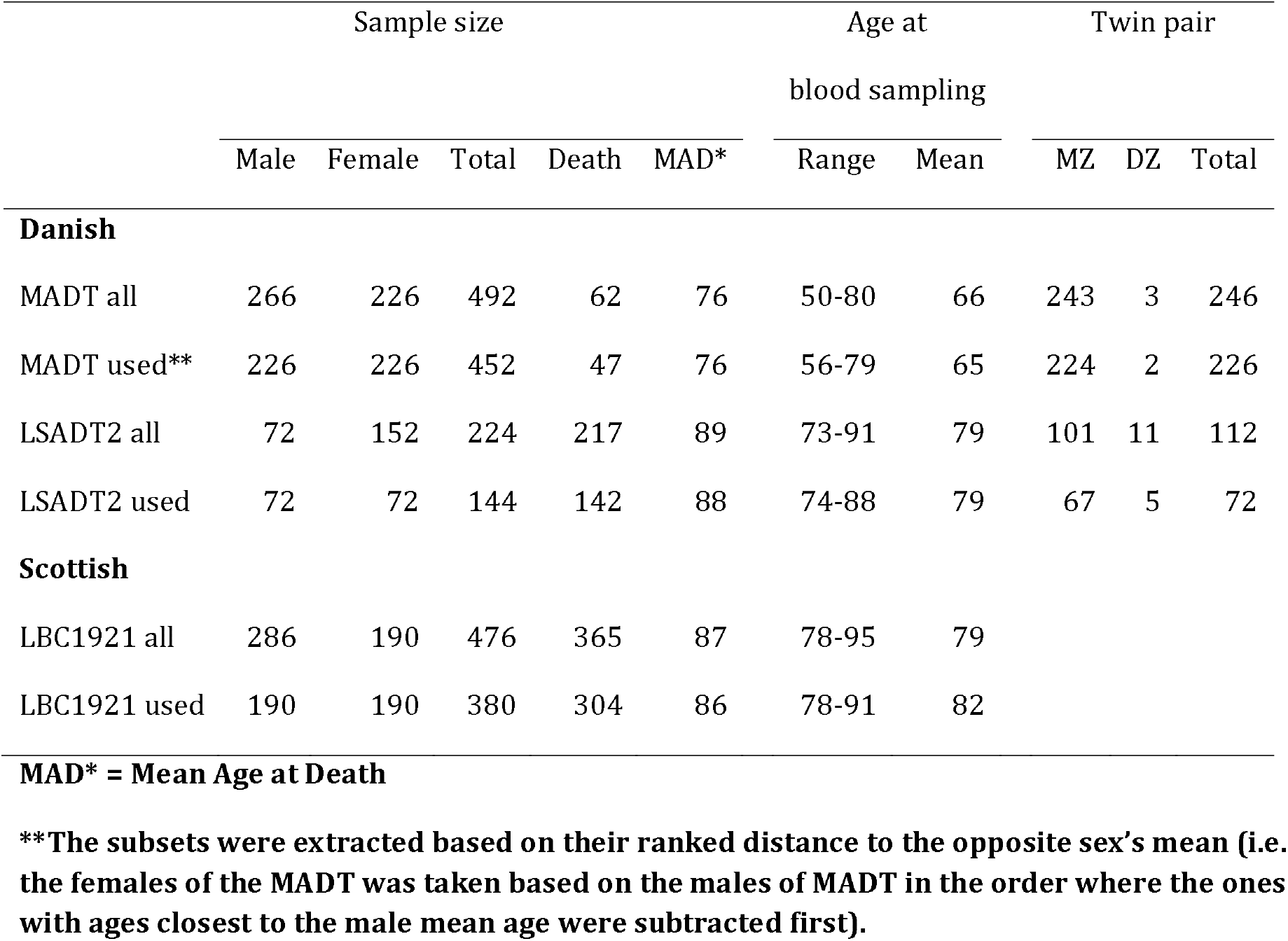
Basic description of samples.

**Table 2.**
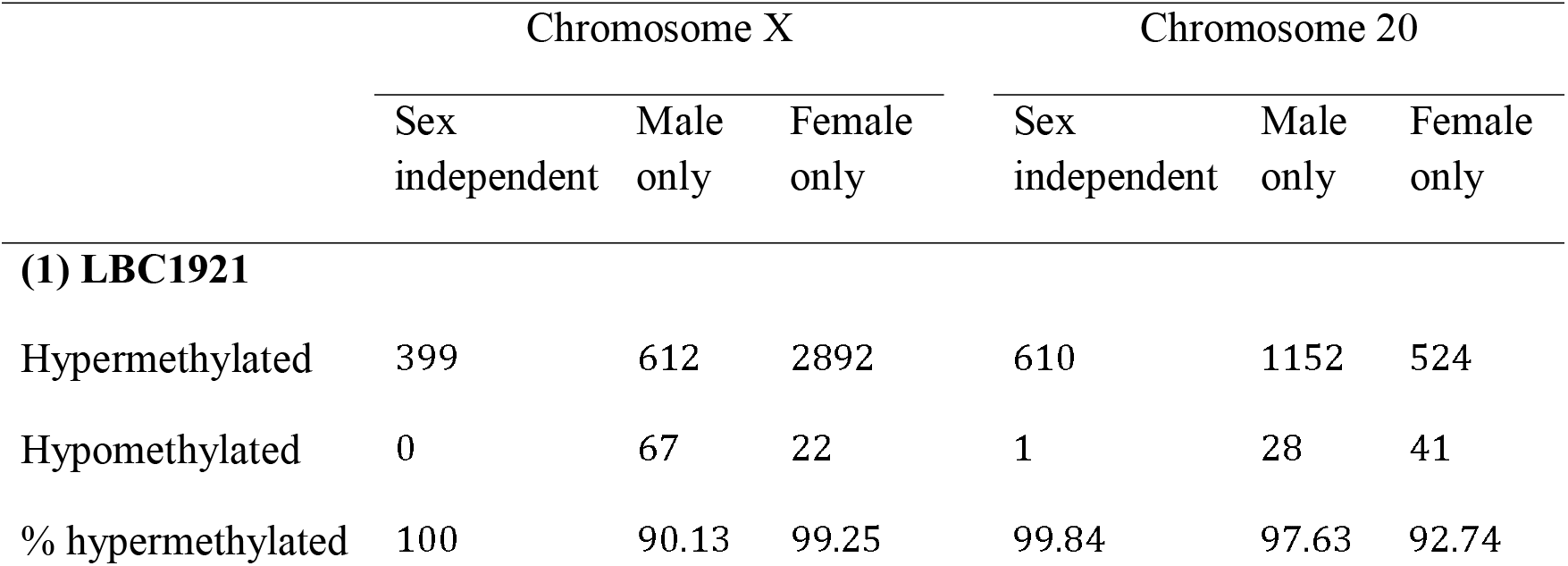
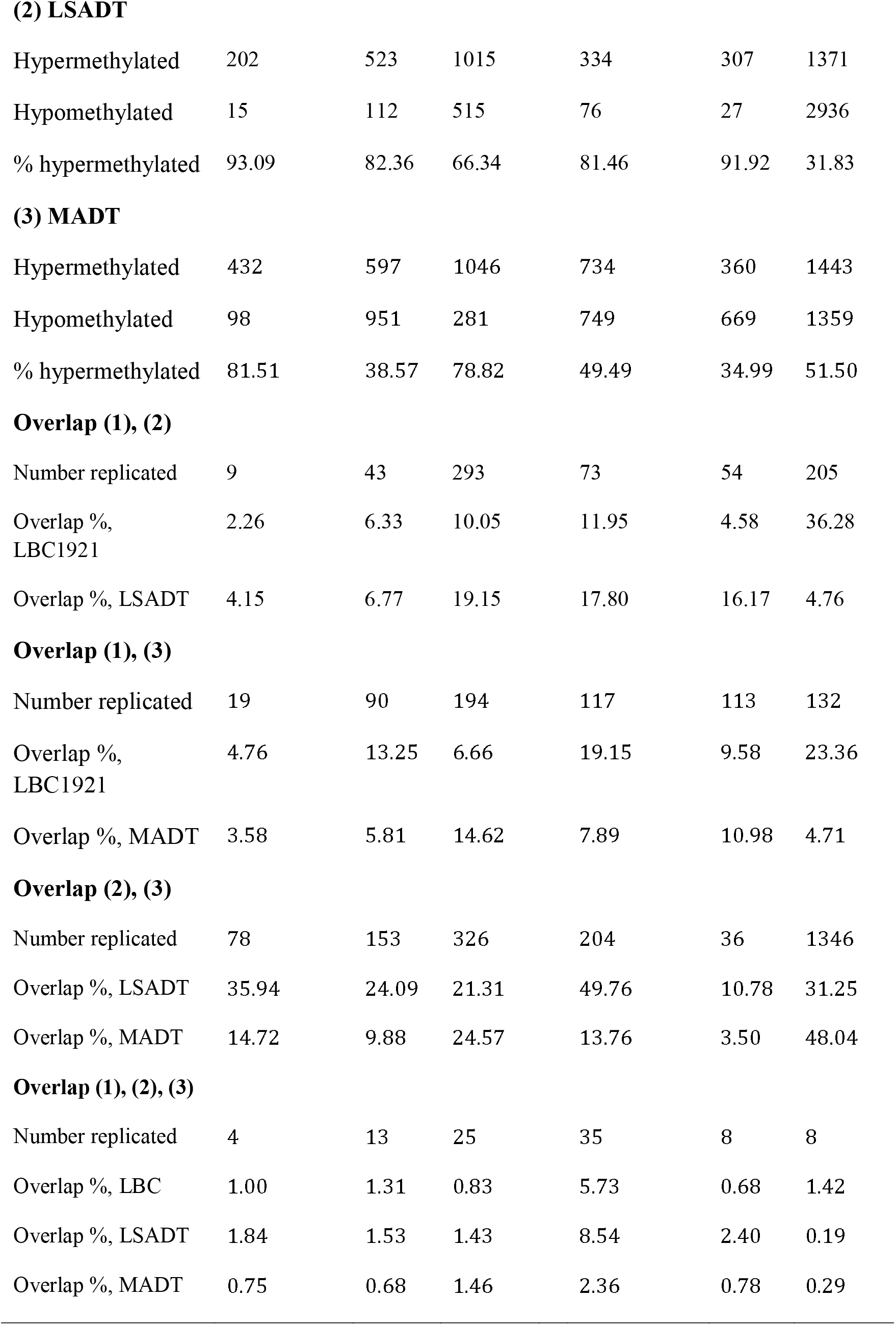
Number of significant sites (FDR<0.05) by cohort and sex with replication results.

|  | Chromosome X |  |  | Chromosome 20 |  |  |
| --- | --- | --- | --- | --- | --- | --- |
|  | Sex<br>independent | Male<br>only | Female<br>only | Sex<br>independent | Male<br>only | Female<br>only |
| **(1) LBC1921** |  |  |  |  |  |  |
| Hypermethylated | 399 | 612 | 2892 | 610 | 1152 | 524 |
| Hypomethylated | 0 | 67 | 22 | 1 | 28 | 41 |
| % hypermethylated | 100 | 90.13 | 99.25 | 99.84 | 97.63 | 92.74 |
| **(2) LSADT** |  |  |  |  |  |  |
| Hypermethylated | 202 | 523 | 1015 | 334 | 307 | 1371 |
| Hypomethylated | 15 | 112 | 515 | 76 | 27 | 2936 |
| % hypermethylated | 93.09 | 82.36 | 66.34 | 81.46 | 91.92 | 31.83 |
| **(3) MADT** |  |  |  |  |  |  |
| Hypermethylated | 432 | 597 | 1046 | 734 | 360 | 1443 |
| Hypomethylated | 98 | 951 | 281 | 749 | 669 | 1359 |
| % hypermethylated | 81.51 | 38.57 | 78.82 | 49.49 | 34.99 | 51.50 |
| **Overlap (1), (2)** |  |  |  |  |  |  |
| Number replicated | 9 | 43 | 293 | 73 | 54 | 205 |
| Overlap %, LBC1921 | 2.26 | 6.33 | 10.05 | 11.95 | 4.58 | 36.28 |
| Overlap %, LSADT | 4.15 | 6.77 | 19.15 | 17.80 | 16.17 | 4.76 |
| **Overlap (1), (3)** |  |  |  |  |  |  |
| Number replicated | 19 | 90 | 194 | 117 | 113 | 132 |
| Overlap %, LBC1921 | 4.76 | 13.25 | 6.66 | 19.15 | 9.58 | 23.36 |
| Overlap %, MADT | 3.58 | 5.81 | 14.62 | 7.89 | 10.98 | 4.71 |
| **Overlap (2), (3)** |  |  |  |  |  |  |
| Number replicated | 78 | 153 | 326 | 204 | 36 | 1346 |
| Overlap %, LSADT | 35.94 | 24.09 | 21.31 | 49.76 | 10.78 | 31.25 |
| Overlap %, MADT | 14.72 | 9.88 | 24.57 | 13.76 | 3.50 | 48.04 |
| **Overlap (1), (2), (3)** |  |  |  |  |  |  |
| Number replicated | 4 | 13 | 25 | 35 | 8 | 8 |
| Overlap %, LBC | 1.00 | 1.31 | 0.83 | 5.73 | 0.68 | 1.42 |
| Overlap %, LSADT | 1.84 | 1.53 | 1.43 | 8.54 | 2.40 | 0.19 |
| Overlap %, MADT | 0.75 | 0.68 | 1.46 | 2.36 | 0.78 | 0.29 |

**Figure 1.**
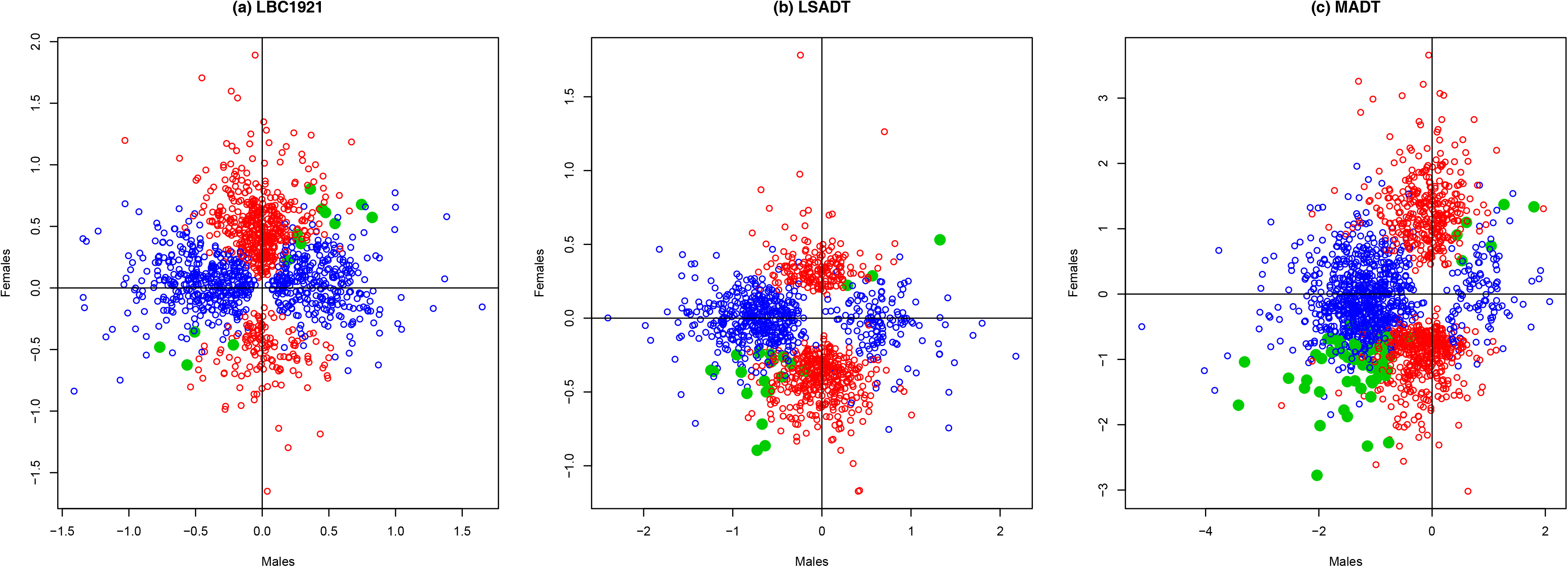
Scatter plots comparing the rate of change in DNA methylation (regression coefficient of age) of significant CpGs (FDR<0.05) between male and female samples for sex-specific (blue for male and red for female only samples) and unspecific (green dots) CpGs on the chromosome X (a, c, e) and chromosome 20 (b, d, f) in LBC1921, LSADT and MADT cohorts.

### Sex-independent age-related methylation patterns

We identified 399, 202 and 432 CpGs hypermethylated, only 0, 15 and 98 CpGs hypomethylated with increasing age in both male and female samples with FDR<0.05 in LBC1921 (Table S1), LSADT (Table S2) and MADT (Table S3) samples, respectively (Table 2, Figure 1). The proportions of hypermethylation were over 81% in all 3 cohorts. Among these sex-independent CpGs, 9 were shared by LBC1921 and LSADT, 19 CpGs by LBC1921 and MADT, 78 by LSADT and MADT (Table 2). The between sample overlapping rates were lower across the 2 countries (<4.76%) than that between the 2 Danish cohorts (>14.72%). Four CpGs overlapped in the 3 cohorts, cg26516882 at 147133598bp, cg15192932 at 153037045bp in the gene body of *PLXNB3*, cg15222604 at 153230625bp in the gene body of *HCFC1*, cg18086582 at 48459799bp in the gene body of *WDR13*.

### Sex-specific age-related methylation patterns

Analysis of the male samples found 612, 523, 597 hypermethylated and 67, 112, 951 hypomethylated CpGs significant (FDR<0.05) only in males of LBC1921 (Table S3), LSADT (Table S4) and MADT (Table S5) cohorts (Table 2, Figure 1). Again, very high proportions of hyper-methylation (>82%) were observed in the older cohorts. In comparison, the proportion was only 39% in the MADT male samples (Figure 1). Among the male-only significant CpGs, 43 overlapped between LBC1921 and LSADT, 90 between LBC1921 and MADT with overlapping proportions <13.25%. In contrast, there were 153 CpGs overlapping between LSADT and MADT male twins with an overlapping proportion of 24.09% for LSADT and 9.88% for MADT (Table 2). Altogether, we found 13 CpGs overlapping across all 3 cohorts (Table 3).

**Table 3.** The 13 age-associated CpGs overlapping in the three samples of males.

| CpG | LBC1921 |  | LSADT |  | MADT |  | Position | Relation to island | RefGene Name | RefGene Group |
| --- | --- | --- | --- | --- | --- | --- | --- | --- | --- | --- |
|  | Coef. | p | Coef. | p | Coef. | p |  |  |  |  |
| cg00817749 | 2.679 | 0.000839 | 38.713 | 5.05E-06 | 18.474 | 4.94E-06 | 37699631 | OpenSea | DYNLT3 | 3'UTR |
| cg01087287 | 1.429 | 0.00111 | 11.703 | 0.004064 | 7.958 | 0.000371 | 134136248 | N_Shore |  |  |
| cg02923300 | 1.584 | 0.001567 | 18.548 | 0.000844 | 6.058 | 0.006026 | 68759322 | S_Shore |  |  |
| cg03199955 | 2.739 | 8.82E-05 | 13.842 | 6.62E-05 | 6.330 | 3.33E-05 | 30671630 | Island | GK | 1stExon;5'UTR |
| cg05070565 | 1.254 | 0.002764 | 23.537 | 1.21E-05 | 6.562 | 0.002654 | 48545185 | OpenSea | WAS | Body |
| cg05587158 | 1.187 | 0.004525 | 16.109 | 0.001506 | 8.647 | 0.000281 | 69487748 | OpenSea | ARR3 | TSS1500 |
| cg11712793 | 3.033 | 1.16E-06 | 19.675 | 0.0013 | 6.693 | 0.003691 | 39548267 | Island |  |  |
| cg12377327 | 4.075 | 3.87E-09 | 27.775 | 4.16E-06 | 8.442 | 0.004333 | 48937697 | Island | WDR45 | TSS200;5'UTR |
| cg13498184 | 1.767 | 0.001832 | 15.158 | 0.000391 | 5.892 | 0.008091 | 144337589 | OpenSea | SPANXN1 | 3'UTR |
| cg15474769 | 1.377 | 0.000698 | 13.832 | 0.002362 | 7.966 | 0.000227 | 53113522 | S_Shore | TSPYL2 | Body |
| cg20521104 | 5.861 | 0.003498 | 35.010 | 1.03E-06 | 14.148 | 0.005475 | 102191578 | N_Shore | RAB40AL | TSS1500 |
| cg24663569 | 1.051 | 0.001176 | 14.530 | 0.002323 | 7.431 | 0.002002 | 70887940 | Island | BCYRN1 | Body |
| cg26028155 | 0.891 | 0.00275 | 18.370 | 3.21E-05 | 6.062 | 0.000967 | 41334113 | Island | NYX | Body |

Analysis of the female samples detected 2892, 1015, 1046 hypermethylated and 22, 515, 281 hypomethylated CpGs significant (FDR<0.05) only in females of LBC1921 (Table S7), LSADT (Table S8) and MADT (Table S9) cohorts (Table 2, Figure 1). As shown by Table 2, many more significant CpGs were found in female than in male samples within each cohort although exactly the same sample sizes were used. Across the cohorts, 293 female-only significant CpGs overlapped between LBC1921 and LSADT, 194 between LBC1921 and MADT, 326 between LSADT and MADT with, in general, higher overlapping proportions than that of the male-only CpGs. Overall, there were 24 CpGs significant only in females in all cohorts with most of them located in the promotors (Table 4). Interestingly, among the genes annotated to the overlapping CpGs in Table 4, *PLXNA3* is also linked to cg15192932 which is one of the four sex-independent CpGs shared by all 3 cohorts.

**Table 4.** The 24 age-associated CpGs overlapping in the three samples of females.

| CpG | LBC1921 |  | LSADT |  | MADT |  | Position | Relation to Island | Gene Name | Gene Group |
| --- | --- | --- | --- | --- | --- | --- | --- | --- | --- | --- |
|  | Coef. | p | Coef. | p | Coef. | p |  |  |  |  |
| cg00011891 | 2.259 | 0.000944 | 5.132 | 0.005058 | 9.864 | 0.003328 | 153714660 | Island | UBL4A | Body |
| cg00705055 | 1.944 | 0.003798 | 7.819 | 0.001572 | 11.487 | 0.000362 | 48824499 | N_Shore | KCND1 | Body |
| cg02033323 | 2.337 | 0.000206 | 5.132 | 0.002027 | 8.173 | 0.001764 | 49118313 | OpenSea | FOXP3 | 5'UTR |
| cg02629737 | 1.454 | 0.003974 | 7.701 | 0.001173 | 7.053 | 0.002951 | 138285540 | Island | FGF13 | 5'UTR;Body |
| cg02763888 | 1.940 | 0.00624 | 4.856 | 0.002699 | 5.294 | 0.004571 | 134427883 | N_Shore | ZNF75D | Body |
| cg04221167 | 1.615 | 0.005207 | 8.473 | 0.000912 | 16.189 | 3.73E-09 | 109683163 | OpenSea | AMMECR1;RGAG1 | 1stExon;5'UTR |
| cg04290766 | 1.811 | 0.010076 | 10.091 | 0.000225 | 13.966 | 3.75E-06 | 152712044 | S_Shore | TREX2 | TSS200 |
| cg05530348 | 2.186 | 0.000663 | 13.253 | 7.93E-05 | 12.001 | 0.003798 | 134558579 | S_Shelf | NCRNA00086 | Body |
| cg05564395 | 1.179 | 0.013364 | 5.167 | 0.000336 | 6.830 | 0.008539 | 48929391 | N_Shore | PRAF2 | 3'UTR |
| cg05600804 | 2.378 | 1.76E-07 | 6.456 | 8.84E-05 | 11.620 | 9.38E-09 | 153623245 | N_Shelf |  |  |
| cg06243722 | 1.355 | 0.009858 | 6.154 | 0.0002 | 10.338 | 6.45E-08 | 3756107 | Island |  |  |
| cg08434396 | 5.162 | 3.85E-05 | 7.594 | 0.001262 | 13.692 | 3.96E-05 | 146993778 | Island | ASFMR1;FMR1 | Body |
| cg08722366 | 3.014 | 0.001046 | 8.234 | 0.006781 | 8.786 | 0.001897 | 153295804 | OpenSea | MECP2 | 3'UTR |
| cg10043957 | 2.608 | 0.009267 | 6.469 | 0.004945 | 18.326 | 1.83E-08 | 40692588 | S_Shore | LOC100132831 | TSS200 |
| cg14002781 | 1.330 | 0.000873 | 2.795 | 0.005482 | 6.342 | 1.65E-06 | 153051367 | OpenSea | IDH3G;IDH3G | 3'UTR;Body |
| cg17536843 | 2.888 | 0.00341 | 6.184 | 0.002077 | 10.619 | 8.93E-05 | 134846527 | OpenSea | CT45A1 | TSS1500 |
| cg18780401 | 4.989 | 4.27E-10 | 4.123 | 0.00787 | 8.677 | 0.001944 | 48815125 | Island | OTUD5 | TSS1500;5'UTR |
| cg21593552 | 4.588 | 0.000151 | 6.312 | 0.000156 | 7.184 | 0.00493 | 135579428 | Island | HTATSF1 | 1stExon;TSS1500;5'UTR |
| cg22265142 | 4.131 | 8.85E-07 | 4.890 | 0.008218 | 7.708 | 0.007261 | 47479101 | Island | SYN1 | 1stExon |
| cg23247655 | 3.965 | 0.001022 | 9.393 | 0.001394 | 9.654 | 0.003695 | 54466701 | OpenSea | TSR2 | TSS200 |
| cg23432159 | 2.900 | 0.002983 | 7.540 | 0.000214 | 9.150 | 0.003921 | 49009222 | N_Shelf |  |  |
| cg23494279 | 1.574 | 0.007304 | 3.874 | 0.006566 | 5.952 | 0.000385 | 153698358 | S_Shore | PLXNA3 | Body |
| cg23731466 | 2.159 | 0.012653 | 5.088 | 0.00198 | 7.498 | 5.44E-05 | 17755254 | Island | SCML1 | TSS1500 |
| cg26101161 | 6.181 | 0.006325 | 15.224 | 3.24E-05 | 23.156 | 2.25E-08 | 122736584 | OpenSea | THOC2 | 3'UTR |

### Relationship between age-related methylation change and mortality

We first fitted Cox models to male and female samples separately. For the male samples, we found 1 (cg10082114, coef.: −0.508, p value: 1.572e-06, FDR: 0.018), 3 (cg09407507, coef.: −1.576, p value: 7.53e-08, FDR: 4.229e-04; cg09703824, coef.: 0.324, p value: 6.81e-07, FDR: 2.55e-03; cg23171231, coef.: −0.267, p value: 6.63e-08, FDR: 4.229e-04) and 72 (Table S10) CpGs associated with mortality in LBC1921, LSADT and MADT cohorts with FDR<0.05. For the female samples, there were 0, 1 (cg05652828, coef.: −0.577, p value: 3.308, FDR: 0.037) and 27 (Table S11) CpGs that were significantly (FDR<0.05) associated with mortality in LBC, LSADT and MADT cohorts. We observed only 1 CpG (cg14728856) significantly correlated with mortality in both males (coef.: −1.979, p value: 4.31e-05, FDR: 0.016) and females (coef.: −2.014, p value: 1.33e-07, FDR: 3.73e-04) of the MADT cohort with nearly equal effect size. The CpG is located in the promotor of *UBA1* gene. In Figure 2, we plot the regression coefficient for each CpG in the Cox model fitted to male data (X-axis) against that for female data (Y-axis) for CpGs with nominal p value <0.05 only in males (blue), only in females (red) and sex-independent (green). The figure shows that the risks of death associated with these CpGs, although mostly not significant after correcting for multiple testing, are predominantly sex-specific, especially in the older LBC and LSADT cohorts.

**Figure 2.**
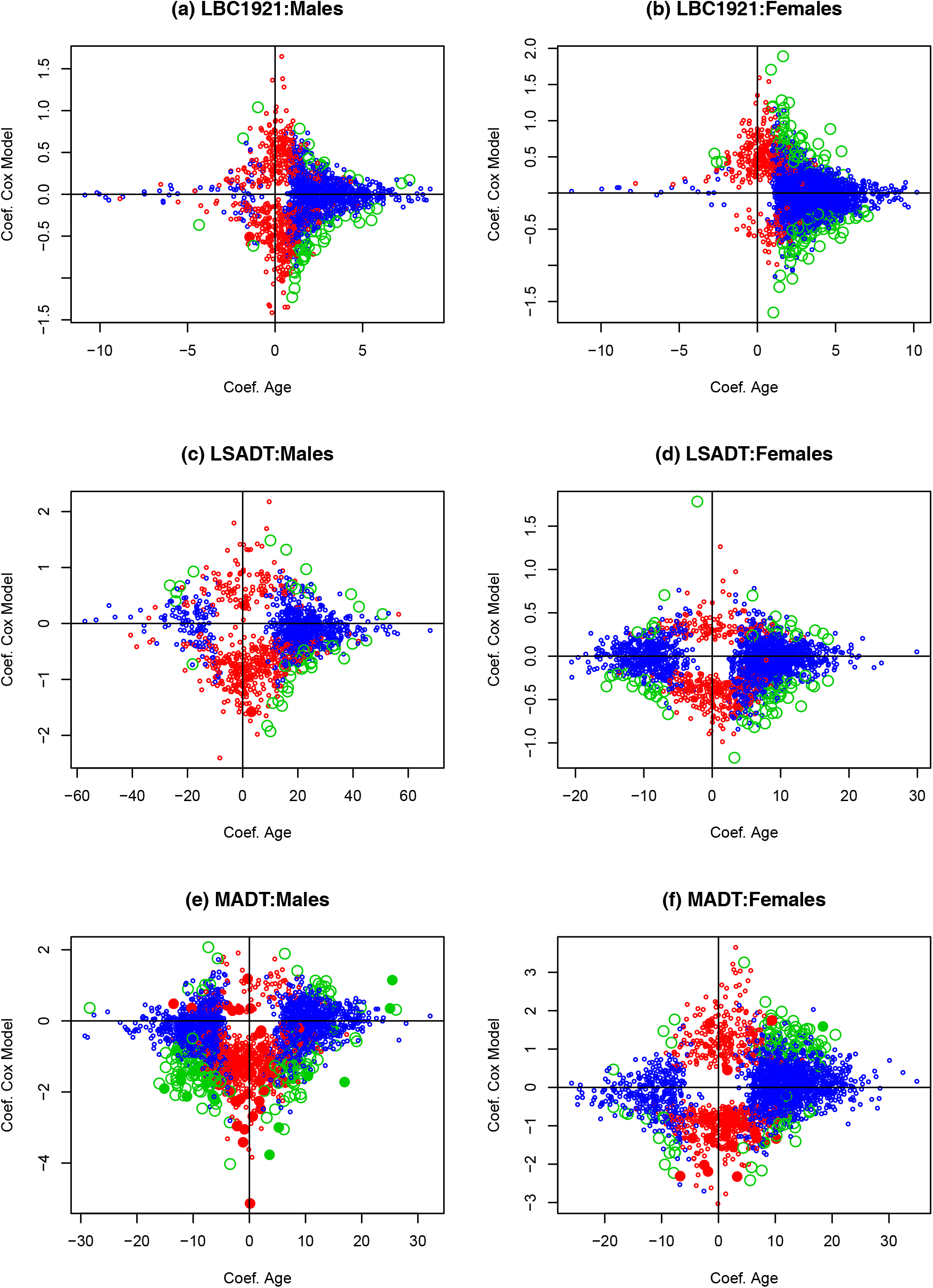
Scatter plots for Cox regression coefficients of CpGs with p<0.05 in male (blue dots) against female (red dots) samples from the LBC1921 (a), LSADT (b) and MADT (c) cohorts. The green dots are CpGs with p<0.05 in both male and female samples.

In Supplementary Table S12, we summarize the relationship between age-associated CpGs (FDR<0.05) and mortality-related CpGs (p<0.05). Interestingly, CpGs hypermethylated with age in males are characterized by high proportion of those with negative Cox coefficients (i.e. reduced risk of death with hazard ration or HR<1) which is significantly different from that of CpGs hypomethylated in aging males. In Females, however, such a consistent pattern was not observed. On the contrary, the female MADT cohort is characterized by a significantly higher proportion of hypermethylated CpGs with HR>1. Figure 3 plots CpGs with FDR<0.05 for age (blue dots) or p<0.05 for mortality (red dots), or both (green dots) in males (left panel) and females (right panel). Additionally, CpGs with FDR<0.05 for mortality are indicated by solid dots. The highly significant mortality-associated CpGs (FDR<0.05) are mostly unrelated to age, except those in solid green in Figures 3e (19 CpGs, Supplementary Table S13) and 3f (2 CpGs, cg02033323, cg04594276) for the younger MADT cohort.

**Figure 3.**
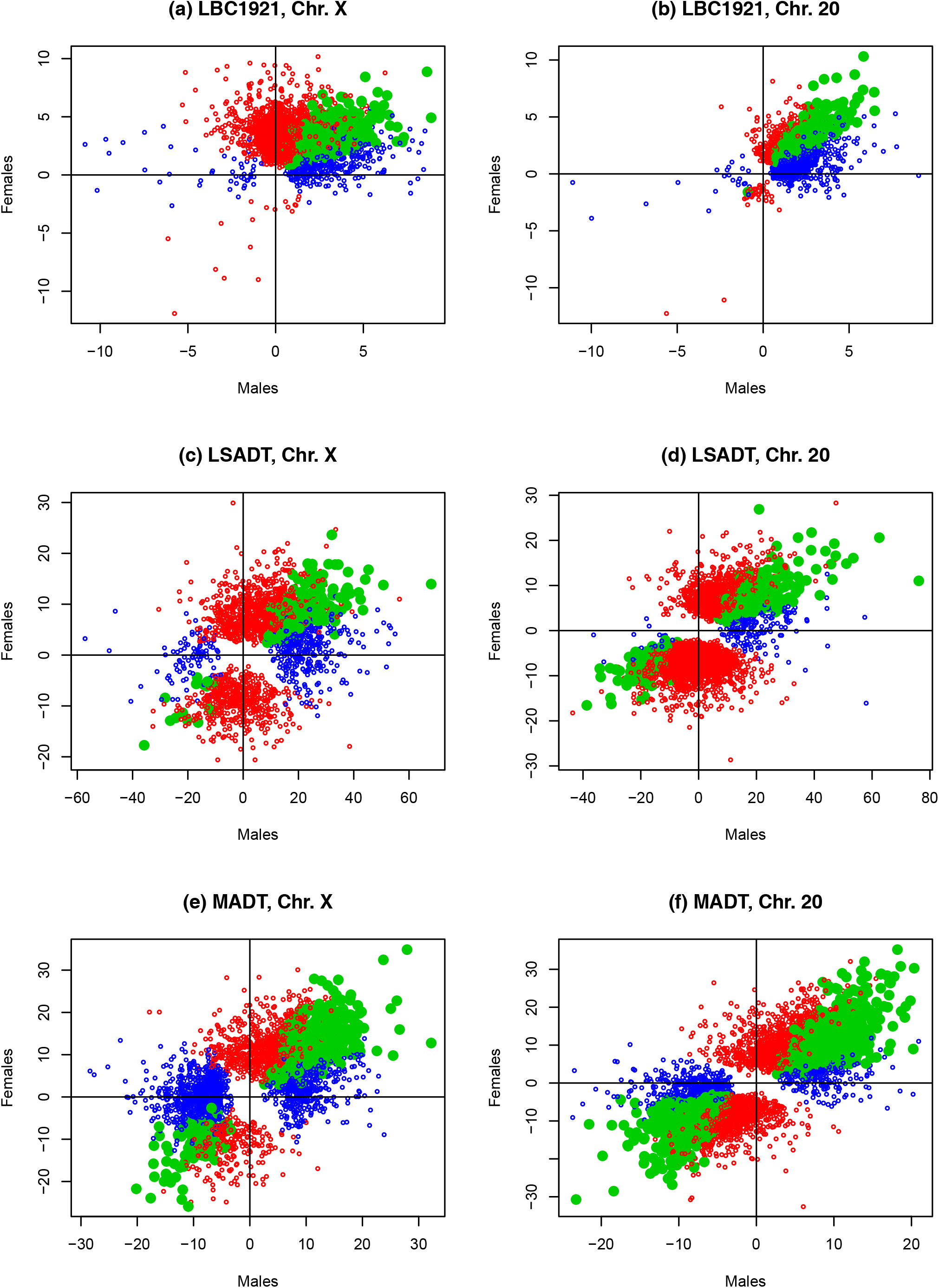
Scatter plots plotting the slope of age for DNA methylation of significant CpGs (FDR<0.05, blue dots) on the X-axis and the effect on mortality (Cox model coefficients for methylation level with p<0.05, red dots) on the Y-axis for the LBC1921 (a, b), LSADT (c, d) and MADT (e, f) cohorts. CpGs with FDR<0.05 for age and p<0.05 for mortality are marked with green colour. CpGs with FDR<0.05 for mortality are indicated as solid dots.

## Discussion

To our knowledge, we have performed the first exploratory analysis of the age-associated DNA methylation patterns on the X-chromosome and their associations with mortality in samples of older people. By estimating the age-dependent change in DNA methylation in male and female samples separately, we were able to discover the sex-specific and non-specific DNA methylation changes across ages and linked these to the risk on mortality while disregarding the sex differences in the levels of X-linked DNA methylation.

As shown in Table 2 and Figure 1, the age-related methylation changes are characterized by a high proportion of hypermethylation except in MADT male samples. The phenomenon could imply that more X-linked sites are methylated with increasing age, or in other words, age-related progressive spreading of hypermethylation on the X-chromosome both in males and in females. Although we deliberately used the same number of samples for males and females, the numbers of significant CpGs varied largely by sex, with many more significant sites found in female (red dots in Figure 1) than in male (blue dots in Figure 1) samples of all 3 cohorts. Since the male and female sample sizes are equal, the observed difference is more likely due to biological reasons rather than statistical artifacts associated with differential power as a result of unequal sample sizes. One could postulate that the X-chromosome could undergo more extensive methylation during aging in females than in males. On the other hand, the phenomenon could also occur if X-linked DNA methylation profiles in males are more variable than in females. More research is needed for clarifying the issue.

For comparison purposes, we performed similar analysis on chromosome 20 which has about same number of CpGs as the X-chromosome measured on the 450K array. In Figure 1, the scatter plots for the coefficients of age estimated in male and female samples do not show the striking difference as compared to the plots for X-Chromosome, except for LBC1921 in Figure 1b where CpGs hypermethylated in females are predominantly also hypermethylated in males or simply became sex-independent (green dots of 610 CpGs). It is interesting to see in Table 2 that, although with about equal number of CpG sites analyzed, we always detected more CpG sites differentially methylated with age in a sex-independent manner on chromosome 20 as compared to X-chromosome. The same pattern applies to the number as well as percentage of sex-independent CpGs overlapping between and across cohorts (Table 2). The patterns could indicate that, in comparison with the autosomal CpGs, the age-dependent methylation changes on the X-linked CpGs are more likely sex-specific.

The sex difference is further observed in the patterns of relationship between age-related methylation and mortality. As shown in Figure 2, although both sexes exhibit a predominant pattern of hypermethylation with age (blue dots), there are significantly more age-associated CpGs that contribute to reduce the risk of death in males (green dots in the down-right of Figure 3a, 3c, 3e). The numbers are clearly shown in Supplementary Table S12 (hypermethylated with age with HR<1). In contrast, a nearly equal number of age-associated CpGs contribute to increase or decrease the risk of death in females. In MADT females, there are even more hypermethylated CpGs with HR>1 (increased mortality with methylation) than HR<1 (decreased mortality with methylation) (Figure 3f, Table S12). The sex-dependent pattern here seems to imply that age-related methylation changes on the X-chromosome are associated with longer male survival but not female, which is contradictory to the observed sex difference in mortality (Owens, 2002). In our recent study, age-related hypermethylation on the Y-chromosome has been shown to lower the risk of death in males (Lund et al. 2019). We think, however, explanation of the phenomenon concerns whether the observed age-related hypermethylation change is the cause or response to aging. In this regard, the longitudinal design that links methylation change at an individual level with risk of death should provide direct estimates of the association and help with causation inference (Li et al. 2018). Together with the result from Lund et al. (2019), we postulate that the observed age-associated hypermethylation on the sex chromosomes in males could mainly represent an active response to the aging process that helps to maintain male survival.

The CpGs in Tables 3 and 4 are those significantly methylated with age in all 3 cohorts in males and females separately. Note that these robustly replicable sex-dependent CpGs are all hypermethylated with increasing age. Among the genes linked to CpGs in Table 4 for females, *PLXNA3* is linked to cg23494279 in the gene body. Interestingly, the gene is also annotated to cg15192932 (gene body), one of the 4 sex-independent CpGs shared by all 3 cohorts. Even more interestingly, the same gene is further among the genes annotated to the 19 age-associated (FDR<0.05) CpGs that also affect mortality (FDR<0.05) in males (Supplementary Table S13, cg01798163 in body of *PLXNA3*). According to the parameter estimates in Table S13, the age-dependent hypermethylation of this CpG is associated with decreased risk of death in males. This gene encodes a class 3 semaphorin receptor and may be involved in cytoskeletal remodeling as well as apoptosis. The gene may be associated with tumor progression and has been found differentially expressed in 15 types of cancer (Saleembhasha & Mishra, 2019). Our results could suggest that activity of one gene can be epigenetically regulated by different models including sex-dependent or independent models through DNA methylation.

In supplementary Table S13, cg17662252 located in the body of *GNL3L* gene is hypermethylated with age with increased risk for death. It has been shown that overexpression of *GNL3L* drives the fraction of genetically defined tumor cells that exhibit markers and tumorigenic properties of tumor initiating cells of enhanced radioresistance and propensity to metastasize (Okamoto et al. 2011). Considering the observation that hypermethylated CpG in the gene body is accompanied by gene overexpression (Yang et al. 2014), our estimate on *GNL3L* is consistent with its role in tumor. Note that the gene is linked to a CpG significant for age and mortality only in males which may contribute to the male–female mortality disparity (Owens, 2002).

In summary, we identified more X-linked CpG sites displaying significant age-related methylation patterns in females than in males, with predominantly high proportions of hypermethylation in the older as compared with the younger cohorts. Survival analysis showed that the age-associated methylation patterns may contribute to reduce mortality in male but not in female samples. Based on our analysis, we conclude that the X-chromosome displays significant age-and sex-dependent methylation patterns which might be differentially associated with mortality in the two sexes. Finally, we point out that, the number of X-linked CpG sites covered by the 450K array is only around 1% of the total number of CpGs on the X-chromosome (∼1.2 million). Considering the limited coverage, interpretation and generalization of our findings in this study should be done with caution. New data collected using high-throughput sequencing techniques such as whole genome bisulfite sequencing should help with replicating and validating our results and verify our conclusions.

## Methods

### The middle-aged Danish twins (MADT)

The MADT samples consist of twin pairs born between 1931 and 1952 collected in the Danish Twin Registry (Gaist et al. 2000). DNA methylation analysis was performed on 492 blood samples (246 monozygotic or MZ twin pairs, 133 male pairs and 113 female pairs) aged from 55 to 80 years with a mean age of 66 (Table 1). For fair cross-sex comparison of significant findings, we took an equal sample size for both sexes as the minimum sample size of the two sexes in statistical analysis. Because there were more male than female samples, this was done by sequentially taking male samples closest to the mean age of female samples until reaching female sample size. This resulted in 113 male and 113 female twin pairs with age range from 56-79 and a mean age of 65 (Table 1).

### The Longitudinal Study of Aging Danish Twins (LSADT)

The LSADT study based on the Danish Twin Registry is a cohort sequential study of elderly Danish twins. LSADT began in 1995 with an assessment of all members of like-sex twin pairs born in Denmark before 1920. Blood samples were drawn during the home visit in 1996 and 1997 from which DNA was isolated and DNA methylation measured recently (Svane et al. 2018). After taking equal sample size for male and female samples, we have in total 144 individuals (72 each sex) included in our analysis with an age range of 74-88 (Table 1). Details on design and data collection were described previously (Christensen *et al.*, 2008).

### The Lothian Birth Cohort 1921 (LBC1921)

The LBC1921 cohort (Deary et al. 2012; Taylor et al. 2018) is a sample of community-dwelling, relatively healthy older people, all of whom were born in 1921. When first recruited at mean age 79 years, most participants were residing in the Lothian region (Edinburgh and its surrounding areas) of Scotland. We use data on the LBC1921 samples collected in the period 1999-2013 from all participants born in 1921. The initial recruitment collected 550 individuals (312 males; 238 females). They have been assessed in five waves, with age ranging from 78 to 95 years. Although the longitudinal study covered multiple waves of follow-up, only the last wave sample from each individual was used to avoid attrition effect in the analysis. As shown in Table 1, after fixing equal sample sizes as did for MADT, we had 190 male and 190 female samples with an age range of 78-91 and a mean age of 82.

### DNA methylation data

DNA methylation profiles for all samples used in this study were analyzed by the same platform, i.e. the Illumina Human Methylation 450K Beadchip containing 485,512 CpG sites across the human genome at single nucleotide resolution in genomic DNA from the whole blood. This study only focused on CpGs from the X-chromosome with a total of 11,232 sites. For comparison purposes, we also included chromosome 20 with 10,383 CpGs (about the same number as the CpGs on X-chromosome) analyzed on the array. Experimental details concerning DNA methylation profiling can be found in Svane et al. (2018) for Danish twins and in Marioni et al. (2015) for the LBC data. The LBC1921 data are accessible through the European Genome-phenome Archive (https://www.ebi.ac.uk/ega/home) with accession number EGAS00001000910.

### Array data preprocessing

For each dataset, normalization on the measured X-chromosome DNA methylation was performed by subset-quantile within array normalization (SWAN) (Maksimovic et al. 2012) using the *minfi* R-package, on male and female X-chromosomes separately. Probes with a detection p-value (a measure of an individual probe’s performance) > 0.01 were treated as missing. CpG sites with more than 5% missing values were removed from the study. After filtering, a total of 11,195 X-linked CpGs were available for subsequent analysis. The similar procedure was applied to chromosome 20 resulting in 10379 CpGs for comparative analysis with the X-chromosome. The normalized methylation data is described as a β value for each site, which is a continuous variable ranging between 0 (no methylation) and 1 (full methylation). Before statistical analysis, the methylation β values were transformed into M values using the logit transformation with M=log_2_(β/(1-β)).

### Correcting for cell type composition

Blood cell-type composition was estimated using Houseman’s methods (based on normalized methylation beta values from 500 differentially methylated regions) for CD8T, CD4T, natural killer cell (NK), B cell, monocyte, and granulocyte using the R packages *minfi* for Danish twins and *celltypes450* (https://github.com/brentp/celltypes450) for the LBC1921s. Correction for cell type composition was done by including the estimated cell type proportions as covariates in the regression models.

### Statistical analysis

For each X-linked CpG site, we fitted a linear mixed effects model to regress its methylation M value on each sample’s age and cell type proportions with array sentrix barcode and sample sentrix position as random effects variables.

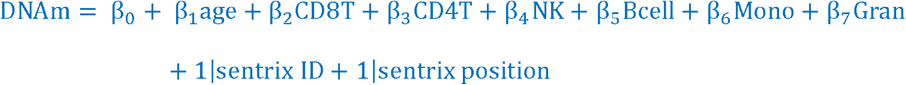

Age-associated change in DNA methylation was assessed by examining the regression coefficient for age β_1_ with β_1_ >0 or β_1_ <0 indicating age-associated hyper-or hypo-methylation. For the Danish twins, the same regression model was fitted but by applying the mixed effect model with the random effect variable defined as twin pair IDs to account for intra-pair correlation on DNA methylation.

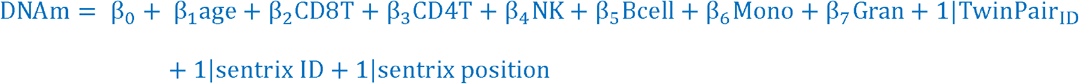

The statistical analysis was done for male and female samples separately to avoid sex differences on the X-chromosome while enabling comparison on sex-specific estimates to identify consistent and sex-dependent age-related DNA methylation patterns. Statistical significance of the CpGs was determined by calculating the false discovery rate (FDR) (Benjamini and Hochberg, 1995) and defined CpGs with FDR<0.05 as significant.

Based on the survival information available for the LBC1921 samples, we additionally fitted Cox regression models to the X-linked CpGs.

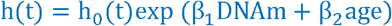

Here, t is the survival time, h(t) is the hazard function, h_0_(t) is the baseline hazard function. The effect of DNA methylation on survival is adjusted for age at methylation measurement. The estimates from the Cox models were used to compare with the estimated slope of age for DNA methylation to examine the relationship between age-dependent methylation and mortality.

## Declarations

### Ethics approval and consent to participate

The MADT study was approved by the Regional Committees on Health Research Ethics for Southern Denmark (S-VF-19980072). The LSADT project has been approved by The Danish National Committee on Biomedical Research Ethics (journal VF 20040241). Written informed consents were obtained from all participants in the Danish twin studies. Ethics permission for the LBC1921 study protocol was obtained from the Multi-Centre Research Ethics Committee for Scotland (MREC/01/0/56) and from Lothian Research Ethics Committee (LREC/2003/2/29).

## Acknowledgments

This work was supported by the Velux Foundation research grant # 000121540. JB is grateful for financial support by VILLUM Young Investigator grant nr. 13154, and H2020 grant REPOTRIAL nr. 777111. We thank the cohort participants and team members who contributed to these studies. Phenotype collection in the Lothian Birth Cohort 1921 was supported by the UK’s Biotechnology and Biological Sciences Research Council (BBSRC), The Royal Society and The Chief Scientist Office of the Scottish Government. Methylation typing was supported by Centre for Cognitive Ageing and Cognitive Epidemiology (Pilot Fund award), Age UK (Disconnected Mind grant), The Wellcome Trust Institutional Strategic Support Fund, The University of Edinburgh, and The University of Queensland. REM and IJD are members of the University of Edinburgh Centre for Cognitive Ageing and Cognitive Epidemiology (CCACE), which is supported by funding from the BBSRC, the Medical Research Council (MRC), and the University of Edinburgh as part of the cross-council Lifelong Health and Wellbeing initiative (MR/K026992/1).

