## Supplementary Table S12 for "Age and mortality associated DNA methylation patterns on the X-chromosome in male and female samples"

Supplementary Table S12. Number of significant sites for age (FDR<0.05) that affect mortality with p<0.05

|  | Males | | |  | Females | | |
| --- | --- | --- | --- | --- | --- | --- | --- |
|  | Hyper | Hypo | Fisher’s Exact Test |  | Hyper | Hypo | Fisher’s Exact Test |
| LBC1921 |  |  |  |  |  |  |  |
| HR>1 | 15 | 2 |  |  | 59 | 3 |  |
| HR<1 | 43 | 2 |  |  | 65 | 0 |  |
| Percent HR>1 | 28.86 | 50.00 | P=0.30 |  | 47.5 | 100 | P=0.113 |
| LSADT |  |  |  |  |  |  |  |
| HR>1 | 10 | 5 |  |  | 14 | 7 |  |
| HR<1 | 35 | 1 |  |  | 39 | 20 |  |
| Percent HR>1 | 22.22 | 83.33 | P=0.006 |  | 26.42 | 25.93 | P=1.00 |
| MADT |  |  |  |  |  |  |  |
| HR>1 | 19 | 7 |  |  | 67 | 4 |  |
| HR<1 | 58 | 107 |  |  | 27 | 15 |  |
| Percent HR>1 | 24.68 | 6.14 | P=0.0007 |  | 71.28 | 20.05 | P=0.0001 |
